# Posture and support geometry, rather than body size, dictate lateral dynamic stability in walking mammalian quadrupeds

**DOI:** 10.64898/2026.06.04.730117

**Authors:** Turgay Akay, Alexander N. Klishko, Claire E. Hanson, Seyed Mohammadali Rahmati, Kenzie G. MacKinnon, Hangue Park, Boris I. Prilutsky

## Abstract

Body size and limb posture vary widely across mammals and are expected to shape locomotor stability, yet direct comparative evidence remains limited. Here, we tested whether smaller, crouched mammals exhibit greater lateral dynamic stability than larger, more upright species by comparing treadmill walking in mice and cats at dynamically similar speeds. Using kinematic analyses and size normalized measures of stability, we show that mice are substantially more laterally stable than cats. This increased stability is associated with relatively wider step widths and more crouched limb posture, indicating that support geometry and posture play dominant roles in stabilizing locomotion. Despite these differences, both species regulate lateral balance on a step-by-step basis, as revealed by relationships between center of mass motion and subsequent adjustments of the border of support. Our findings demonstrate that locomotor stability does not scale simply with body size but depends critically on posture dependent strategies that differ across species. These results identify lateral stability as a key factor of locomotor adaptation and suggest that crouched postures in small mammals may reduce reliance on active neural control while enhancing robustness in complex environments.

**SUMMARY STATEMENT:** Lateral dynamic stability during quadrupedal locomotion depends primarily on limb posture and support geometry rather than body size. Smaller mammals achieve greater stability through crouched postures and wider step widths, whereas larger mammals operate closer to stability limits and rely more heavily on active control.

## INTRODUCTION

Mammalian locomotion exhibits pronounced diversity in limb posture and balance control, reflecting critical adaptations to body size and environmental demands (Gray, 1968; Gambaryan, 1974; Nevo, 1979; Biewener, 1989; Hildebrand, 1989; Fischer and Blickhan, 2006). Larger mammals typically employ extended limb postures that optimize musculoskeletal stress, energetic efficiency and speed but may compromise lateral stability (Gambaryan, 1974; Alexander and Jayes, 1983; Biewener, 1989). Conversely, small mammals like mice utilize metabolically expensive crouched postures (Taylor et al., 1982; Horner et al., 2016; Thirkell et al., 2025) that likely enhance maneuverability and stability, potentially compensating for their slow locomotor speed and thus vulnerability to predation (Alexander, 2002; Walter, 2003; Fuentes, 2016; Riskin et al., 2016; Dick and Clemente, 2017). However, body size further complicates stability via competing mechanical and neural factors: larger animals experience longer sensorimotor delays but possess greater body inertia that passively resists destabilizing forces (More and Donelan, 2018; Mohamed Thangal and Donelan, 2020). It remains unresolved how these opposing biomechanical effects interact with limb posture to shape lateral stability across species. Resolving this gap is particularly important given the widespread use of both small rodents and larger mammals to study the neural control of human locomotion and balance (Vidal et al., 2004; Frigon, 2020; Latash et al., 2020; Santuz et al., 2022; Modi et al., 2024).

To effectively quantify lateral stability during locomotion, the dynamic margin of stability (*MoS*) is widely used, as it incorporates both center of mass (*CoM*) position and velocity relative to the base of support (Hof et al., 2005; Gates et al., 2013; Park et al., 2019; Nguyen et al., 2023). While a wider step width increases static stability, it can paradoxically amplify the destabilizing effects of lateral *CoM* motion (Farrell et al., 2014; Molina et al., 2023).

Here, we aimed to separate the contributions of animal size and limb posture to dynamic stability by comparing basic limb kinematics, lateral *CoM* dynamics, and a size-normalized margin of stability, *MoS/l_0_* (Nguyen et al., 2023), between the domestic cat and the wild-type mouse. We hypothesized that size-independent morphological features, such as relative step width and the degree of crouching, dictate the pronounced differences in dynamic stability between these distinct species.

Beyond baseline stability, we also investigated the active strategies these species use to maintain lateral balance. Bipedal humans correct lateral *CoM* deviations via step-by-step modulation of foot placement (Townsend, 1985; Bauby and Kuo, 2000; Rankin et al., 2014; Wang and Srinivasan, 2014). While recent work suggests similar principles may operate in forelimbs during hexapedal or diagonal trotting (De Comite and Seethapathi, 2025), lateral forefoot placement may be less effective during typical quadrupedal walking because a forelimb contacts the ground while the ipsilateral hindlimb is already in stance.

We therefore hypothesized that quadrupeds (specifically cats and mice) maintain lateral stabilization through two distinct neuromotor control strategies during walking: (1) modulation of the swing onset timing of the contralateral forelimb to adjust the onset of the ipsilateral double-support phase and/or (2) actively adjusting the lateral border of support—center of pressure (*CoP*) during this same double-support phase—by appropriate placement of the ipsilateral forepaw on the ground. During the ipsilateral double-support, the gravitational force vector acting on the center of mass (*CoM*) generates a moment of force in the frontal plane. This moment acts as a mechanical brake, decelerating and reversing lateral body motion, offering a shared, robust mechanism for step-by-step stabilization across mammalian quadrupedal walking (Park et al., 2019; Latash et al., 2020).

## MATERIALS AND METHODS

### Animals

Experiments were conducted on four adult female cats (mass 2.55–4.11 kg, age 1–2 years) and five adult wild-type mice (3 female, 2 male; mass 18.4–28.1 g; age 67–91 days; see **Table 1**). All procedures involving cats complied with the US Public Health Service Policy on Humane Care and Use of Laboratory Animals and were reviewed and approved by the Institutional Animal Care and Use Committee of the Georgia Institute of Technology (protocol #A100011). All procedures involving mice followed the guidelines of the Canadian Council on Animal Care and were approved by the University Committee on Laboratory Animals at Dalhousie University (protocol #21-109).

**Table 1.** Animal characteristics.

| Cats | Sex | Mass,<br>kg | Thigh<br>length,<br>cm | Shank<br>length,<br>cm | Tarsals<br>length,<br>cm | Total<br>hindlimb<br>length,<br>cm | Cycles<br>analyzed |
| --- | --- | --- | --- | --- | --- | --- | --- |
| MO | F | 4.11 | 10.7 | 12.5 | 6.9 | 30.1 | 37 |
| NO | F | 3.91 | 9.5 | 11.0 | 6.7 | 27.2 | 21 |
| TA | F | 3.00 | 10.1 | 10.2 | 6.4 | 26.7 | 48 |
| WE | F | 2.55 | 9.7 | 10.1 | 5.9 | 25.7 | 22 |
| Mean±SD |  | 3.39±0.74 | 10.0±0.5 | 11.0±1.1 | 6.5±0.4 | 27.4±1.9 |  |
| Mice |  | g |  |  |  |  |  |
| M1 | M | 24.6 | 1.5 | 1.6 | 0.9 | 4.0 | 24 |
| M2 | F | 21.8 | 1.4 | 1.5 | 0.8 | 3.7 | 13 |
| M3 | F | 18.4 | 1.4 | 1.4 | 0.8 | 3.6 | 4 |
| M4 | F | 20.3 | 1.3 | 1.5 | 0.9 | 3.7 | 15 |
| M5 | M | 28.1 | 1.6 | 1.7 | 0.8 | 4.1 | 2 |
| Mean±SD |  | 22.6±3.8 | 1.4±0.1 | 1.5±0.1 | 0.8±0.1 | 3.8±0.2 |  |

In accordance with the ARRIVE 2.0 guidelines for animal research (Percie du Sert et al., 2020), we minimized animal use by employing the same animals across multiple studies addressing distinct scientific questions. The cats included in the present study were also used in previous investigations (Park et al., 2019; Latash et al., 2020; Klishko et al., 2021; Prilutsky et al., 2022).

### Electrode implantation surgeries

Although electromyographic (EMG) recordings are not reported in the present study, all cats and mice underwent EMG implantation surgeries as part of other projects.

*Cats*. Before implantation surgeries and kinematic data collection, cats were trained using positive reinforcement to walk on a split-belt treadmill at tied-belt speeds ranging from 0.4 to 0.8 m/s and to tolerate placement of 28 reflective markers on shaved skin (Park et al., 2019). EMG electrodes were implanted into major muscles of the right hindlimb under general anesthesia and aseptic conditions, following procedures described previously (Prilutsky et al., 2011; Klishko et al., 2021). After surgery, animals recovered for two weeks. Analgesia was provided for three days using a fentanyl transdermal patch (2–25 μg/h) and/or subcutaneous injection (s.c.) of buprenorphine (0.01 mg/kg) or ketoprofen (2 mg/kg). Antibiotics were administered for ten days using cefovecin (8 mg/kg, s.c.) or ceftiofur (4 mg/kg, s.c.).

*Mice.* Mice were not trained prior to surgeries or experiments. Eight bipolar EMG electrodes (Pearson et al., 2005) were implanted into right hindlimb muscles under general anesthesia, following procedures described previously (Akay et al., 2006; Santuz et al., 2022). Postoperative analgesia consisted of buprenorphine (0.03 mg/kg, s.c.) and ketoprofen (5.0 mg/kg, s.c.). After surgery, mice were housed in a heated recovery cage for three days and then returned to their home cages. Food mash and hydrogel were provided during the first three days of recovery. Additional buprenorphine injections were administered at 12-hour intervals for 48 hours. Locomotor recordings began after full recovery, at least ten days after surgery.

Although EMG activity was recorded during locomotion throughout the experimental period as part of related projects, these data are not reported in the present study.

### Behavioral recording sessions

Because experiments on cats and mice were conducted in two different laboratories, procedures for each species are described separately below.

*Cats.* Three-dimensional whole-body kinematics of walking cats were recorded at 120 Hz using a six-camera motion-capture system (Vicon Motion Systems Ltd., UK). Small reflective markers were attached to shaved skin overlying major hindlimb and forelimb joints, as well as bony landmarks on the digits, pelvis, scapulae, and head (Farrell et al., 2014; Park et al., 2019). Prior to kinematic analysis, digitized marker trajectories were low-pass filtered using a fourth-order, zero-lag Butterworth filter with a cutoff frequency of 6 Hz.

Hindlimb and forelimb segment lengths were measured for each cat using an anthropometer while the animal was sedated (dexmedetomidine, 40–60 μg/kg, intramuscular injection) prior to shaving. Segment lengths and measured body mass (**Table 1**) were used to estimate segment mass and relative center-of-mass locations based on regression equations developed in (Hoy and Zernicke, 1985). Using these segment parameters and filtered marker coordinates, we computed center-of-mass positions for each body segment and the whole-body *CoM* (Farrell et al., 2014; Park et al., 2019); **Figure 1A**.

**Figure 1.**
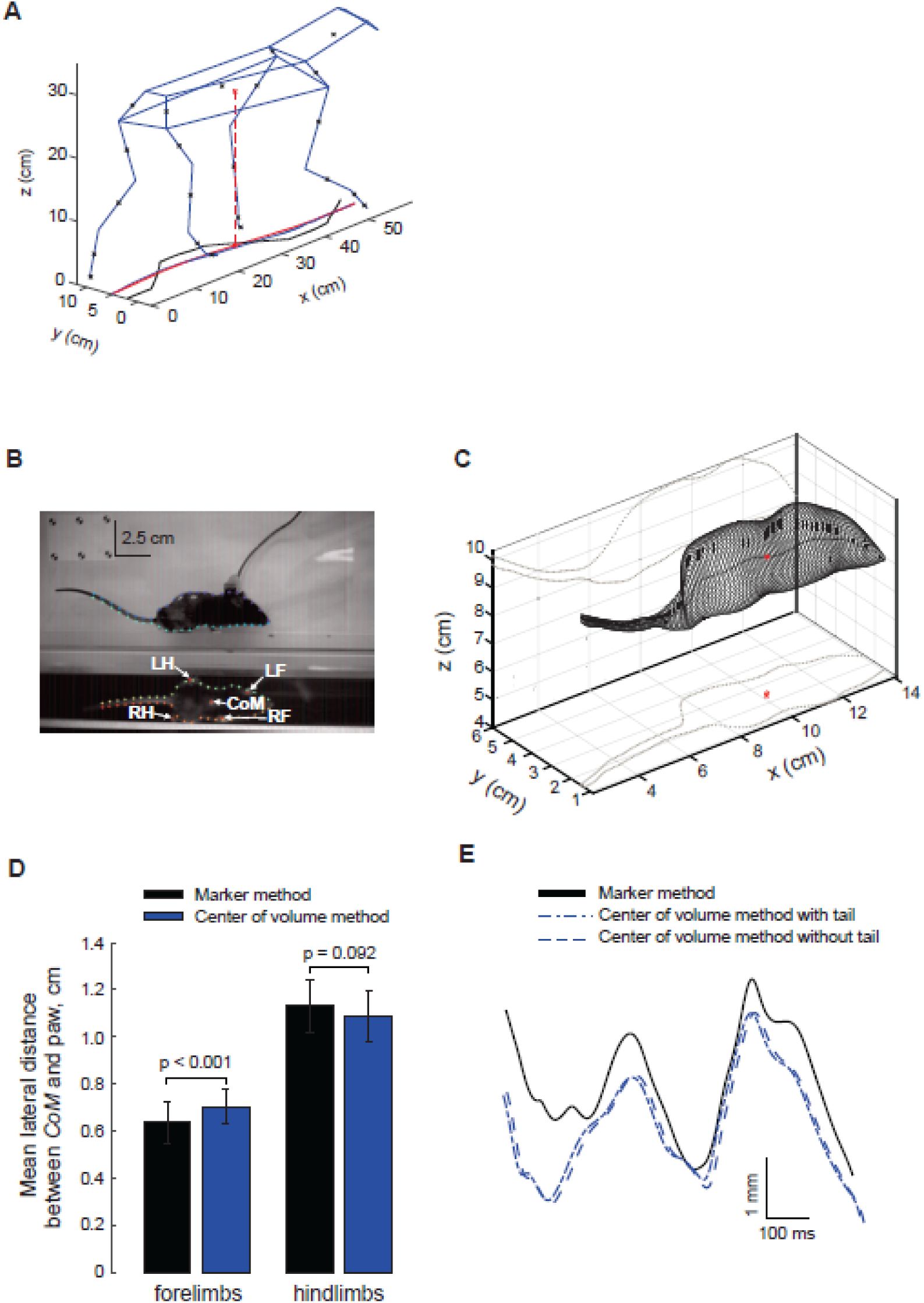
Determination of the center of mass (*CoM*) displacement during locomotion of cats and mice. **A**: Illustration of a method to determine cat lateral *CoM* position from 3D motion capture and individual segment *CoM* positions (black crosses). Body *CoM* position and its projection on the transverse plane are shown by red crosses. Also shown on the transverse plane are schematics of trajectories of the *CoM* (red line), extrapolated *CoM* (*xCoM*, blue line), and center of pressure (*CoP*, black line). **B-E:** Illustration of two methods to determine lateral body *CoM* position in walking mice. In the first method, we painted a marker just caudal to the tip of the sternum on the ventral side of the body corresponding to the *CoM* projection on the mouse transverse plane. This marker and mouse paws are shown on the transverse mouse image in **B** (CoM, LH, LF, RH and RF indicate painted markers on the *CoM* projection and paws of left hindlimb, right forelimb, right hindlimb and right forelimb, respectively). In the second method, we automatically digitized contours of the sagittal and transverse mouse images using a machine learning software (circles marking the contours) after the machine learning program was trained using manual digitization of selected trials (crosses in **B**). Digitized contour points in the sagittal and transverse images were fitted by ellipses, and the center of volume of 3D body shape, corresponding to the *CoM*, was computed (**C**, red cross). Red circle and red cross in the transverse plane in **C** correspond to vertical projections of the *CoM* determined by the two methods. **D**: The mean (±SD) lateral distance between *CoM* and hindpaws and forepaws in a walking cycle of a mouse determined by the two methods. The difference in distance between the two methods was significant, but small for forelimbs and non-significant for hindlimbs (the paired samples t-test, n = 10). **E**: An example of mouse *CoM* trajectories determined by the marker method, by the center of volume method with inclusion of tail in the mouse volume, and by the center of volume method without the tail.

*Mice.* Kinematic data were collected from mice walking on a custom-built treadmill with a transparent belt and an angled mirror positioned at 45° beneath the belt, allowing simultaneous acquisition of sagittal-and horizontal-plane high-speed video recordings at 500 frames per second (Santuz et al., 2022). We used DeepLabCut (DLC) v2.1.7 (Mathis et al., 2018) to obtain sagittal-plane coordinates of painted right hindlimb markers on the metatarsophalangeal (MTP), ankle, knee, and hip joints, while horizontal-plane coordinates were obtained for all four paws and for the projection of the body *CoM*, marked by a white dot on the ventral surface of the trunk (**Figure 1B**).

The longitudinal position of the *CoM* along the body axis was determined in five mice in pilot experiments. Each mouse was euthanized and frozen overnight at −20 °C in a posture approximating the walking posture. Once rigid, the longitudinal *CoM* location was identified by balancing the body to determine the point of static equilibrium. In all five mice, this point was located just caudal to the tip of the sternum.

Digitized horizontal-plane coordinates of the *CoM* and paw positions, as well as sagittal-plane coordinates of right hindlimb joints, were transferred to Spike2 (Cambridge Electronic Design, UK) for further analysis using custom-written code in the Spike2 programming environment.

We validated the method of estimating the *CoM* position from the ventral marker by independently computing the center of body volume based on digitization of mouse body contours in sagittal and transverse views using an overlaid grid (**Figure 1B**). This validation was performed in three mice walking overground on the same treadmill with the belt stationery, as the moving transparent belt reduced the visibility of body contours. Assuming uniform density of the mouse body, the center of body volume coincides with the *CoM*.

Ten representative overground walking trials from three mice were manually digitized (crosses in **Figure 1B**) and used to train DeepLabCut through more than 500,000 iterations to automatically digitize unlabeled grid–contour intersection points in 70 videos from five mice walking overground (circles in **Figure 1B**). Ellipses were fitted to the corresponding contour points to reconstruct the three-dimensional body shape, from which the center of body volume was computed (**Figure 1C**).

To compare the two *CoM* estimation approaches, we calculated the mean lateral distances between the *CoM* position and the hindpaw and forepaw positions obtained using each method. The average differences between methods were 9.4% for hindpaws and 4.1% for forepaws (**Figure 1D**). Excluding the tail from the calculations did not substantially affect the estimated *CoM* position (**Figure 1E**). Because both approaches yielded similar estimates and the center-of-volume method could not be applied to treadmill recordings, we used the ventral *CoM* marker-tracking method for all analyses of mouse treadmill locomotion in this study.

### Data analysis

We compared locomotor kinematics and lateral dynamic stability between cats and wild-type mice during treadmill walking at speeds of 0.4 m/s for cats and 0.1 m/s for mice. These speeds correspond to comparable Froude numbers (Fr = 0.065 for cats and Fr = 0.05 for mice). The Froude number is defined as 𝐹𝑟 = 𝑣^2^/𝑔 · 𝑙_ℎ𝑖𝑝_ (where 𝑣 is locomotion speed in m/s; 𝑙_ℎ𝑖𝑝_ is the mean hip height during stance (approximately 0.25 m for cats and 0.02 m for mice), and 𝑔 = 9.81 m/s^2^ is gravitational acceleration. Because the Froude number represents a size-normalized measure of locomotion speed, it allows comparison of kinematic and stability variables across species under the assumption of dynamic similarity (Alexander and Jayes, 1983).

We quantified major kinematic variables of walking, including cycle duration, stance duration, duty factor (stance duration divided by cycle duration), joint angles, and the length and orientation of the right hindlimb (RH) during walking. Hip angle definitions differed between cats and mice because a marker on the iliac crest was not available in mice (see joint-angle definitions in **Figure 2A**). RH length during walking was defined as the distance between the hip and metatarsophalangeal (MTP) joints (𝑙_ℎ𝑖𝑝−𝑀𝑇𝑃_) and RH orientation was defined as the angle between the horizontal and the line connecting the hip and MTP joints (Klishko et al., 2014); **Figure 2A**. Maximal and minimal values of RH orientation corresponded to swing onset (cycle onset) and stance onset, respectively (Pantall et al., 2012). Coordinates of the right knee joint were corrected using measured thigh and shank lengths together with recorded hip and ankle coordinates, following established methods (Goslow et al., 1973).

**Figure 2.**
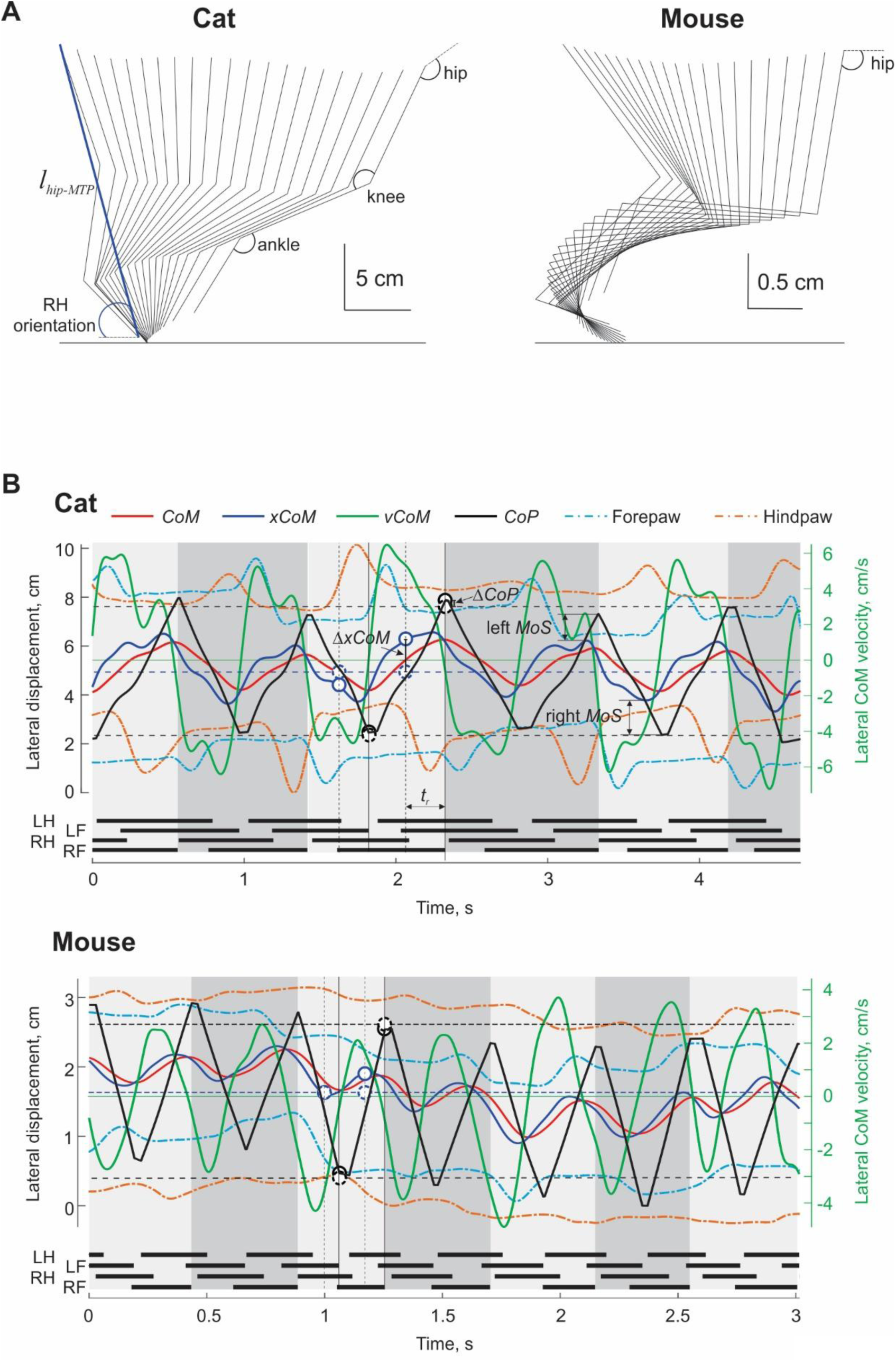
Characteristics of locomotor kinematics and lateral stability in mice and cats. **A**: Stick figures of right hindlimb (RH) metatarsals, skank and thigh during the stance phase of walking. Ankle and knee joint angles are defined in both species as shown in the left panel. The hip joint angle in cats is defined as the angle between the thigh and the line connecting the big trochanter and the iliac crest (not shown); in mice, the hip joint angle is defined as the angle between the thigh and the horizontal. The left panel shows the definition of the RH functional length (the distance between the hip and metatarsophalangeal (MTP) joints, 𝑙_ℎ𝑖𝑝−𝑀𝑇𝑃_) and hindlimb orientation (RH orientation). **B**: Examples of lateral displacements of the center of mass (*CoM*, red continuous lines), extrapolated center of mass (*xCoM*, blue continuous lines), center of pressure (*CoP*, black continuous lines), and forepaw and hindpaw (orange and cyan dash-dot lines, respectively) and the center of mass lateral velocity (*vCoM*, green continuous lines) during walking in a cat (top) and a mouse (bottom). Thick horizontal black lines below the *CoM* and paw kinematic traces represent the stance phases of the left and right forelimbs (LF and RF) and left and right hindlimbs (LH and RH). Left *MoS* and right *MoS* are left and right margines of dynamic stability. Δ𝑥𝐶𝑜𝑀 is a deviation of *xCoM* lateral position (blue continuous circles) at the middle of a diagonal double support or quadrupedal support phase (vertical black dashed lines) from the mean *CoP* (blue dashed circles on the horizontal blue dashed line) calculated across all strides within a trial. Δ𝑥𝐶𝑜𝑃 is a deviation of the *CoP* at the onset of the ipsilateral double support phase (black continuous circles) from the mean ipsilateral *CoP* peaks (black dashed circle on the horizontal black dashed line) calculated across all strides within a trial.

To quantify lateral stability during walking, we computed hindlimb and forelimb step width as the difference in lateral position between the corresponding left and right paws at stance onset. Lateral dynamic stability was quantified using the margin of stability (*MoS*), defined as the distance between the peak position of the extrapolated center of mass (*xCoM*) and the center of pressure (*CoP*) at the onset of the ipsilateral double-support phase in each stride (Park et al., 2019); **Figure 2B** (left *MoS* and right *MoS*). *MoS* values computed for the left and right sides within each stride were averaged.

The *xCoM* was calculated as (Hof et al., 2005): 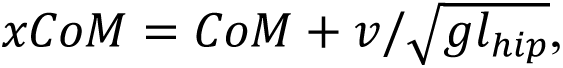 where 𝐶𝑜𝑀 and 𝑣 are the lateral position and velocity of body *CoM*, 𝑙_ℎ𝑖𝑝_ is the mean hip height during the stance phase, and 𝑔 is gravitational acceleration. The *CoP* was computed as 𝐶𝑜𝑃 = (𝑦_𝐿_𝐹_𝐿_ + 𝑦_𝑅_𝐹_𝑅_)/(𝐹_𝐿_ + 𝐹_𝑅_), where 𝑦_𝐿_ and 𝑦_𝑅_ are the mean lateral positions of the left and right limb paws (forelimb and hindlimb), respectively, and 𝐹_𝐿_ and 𝐹_𝑅_are the corresponding resultant vertical ground-reaction forces. As demonstrated previously – see Figure 2 in (Park et al., 2019), these forces can be accurately approximated from body weight and the timing of the ipsilateral double-support phases, during which only the ipsilateral forelimb and hindlimb support the body. Such phases were consistently present in cats walking at 0.4 m/s, a typical low walking speed for cats.

Mice walked intermittently at 0.1 m/s but more frequently adopted a trotting gait. To enable direct comparison of stability between species within the same gait, analyses were restricted to walking cycles only (**Figure 2B**). Only during walking do ipsilateral double-support phases occur, allowing 𝐹_𝐿_and 𝐹_𝑅_to be approximated as body weight and enabling accurate computation of *CoP*.

All translational kinematic variables were normalized by the total anatomical length of the right hindlimb (𝑙_𝑅𝐻_), defined as the sum of thigh, shank, and tarsal segment lengths for each animal (**Table 1**), to account for differences in body size. The mean height of the right hip during the stance phase, normalized by 𝑙_𝑅𝐻_, was used to quantify the degree of crouching during locomotion (Gatesy and Biewener, 1991; Riskin et al., 2016). Normalized lateral *CoM* velocity was computed as the square root of the Froude number: 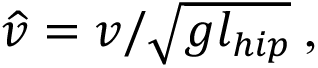 where 𝑣 is the lateral *CoM* velocity in m/s. Normalized stride cycle duration was computed as (Hof, 2018): 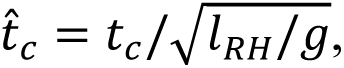 where 𝑡_𝑐_ is the stride cycle duration in seconds.

We further normalized the average margin of stability (*MoS*) in each stride, as well as some other stability related measures (e.g., stance width, stride length), by the total anatomical length of the right hindlimb 𝑙_𝑅𝐻_and by the mean hip-MTP distance 𝑙_ℎ𝑖𝑝−𝑀𝑇𝑃_during the stance phase. Normalization by 𝑙_𝑅𝐻_was used to obtain the measure of lateral dynamic stability that is independent of overall animal size (Nguyen et al., 2023). Normalization by the mean 𝑙_ℎ𝑖𝑝−𝑀𝑇𝑃_distance accounts for limb posture, as locomotor stability depends on both body size (height) and limb configuration (Alexander, 2002; Hof et al., 2005). Comparing stability measures normalized by these two methods therefore allowed us to separate the contributions of animal size and limb posture to differences in dynamic stability.

Normalizing *MoS* by a length scale yields a dimensionless measure that can be interpreted as the impulse of force, applied at the *CoM*, required to destabilize the body, independent of body mass and linear size (Hof et al., 2005; Nguyen et al., 2023). The corresponding non-normalized impulse of force that causes the *xCoM* to cross the boundary of support when applied to the *CoM* at the instant of the *xCoM* peak is given by: 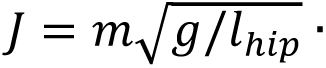 where 𝑚 is body mass (Hof et al., 2005). Assuming that this impulse is applied over the duration of the ipsilateral double-support phase (𝑡_𝑖𝑑𝑠𝑝_), we computed the corresponding destabilizing lateral force acting on the *CoM* as 𝐹_𝑙𝑎𝑡_ = 𝐽/𝑡_𝑖𝑑𝑠𝑝_.

We also investigated whether/how cats and mice correct their lateral *CoM* motion within a walking cycle. We defined the state of *CoM* motion towards a lateral border of support by the lateral deviation of the *xCoM* (blue continuous circles in **Figure 2B**) from the mean *CoP* position computed across all strides within a given trial (blue dashed circles on the horizontal blue dashed line) at the middle of the diagonal double support phase (or quadrupedal support phase at slower strides); this instant is indicated by the vertical dashed lines (**Figure 2B)**. The specific instant for evaluation of the *COM* motion state was selected because at this time point the lateral *CoM* position is at or close to the mid distance between the left and right borders of support on average, and the lateral *CoM* velocity towards the border of support is near its peak (**Figure 2B**).

We assumed that when *xCoM* substantially deviated towards a lateral border of support, the animal would adjust the limb support configuration to prevent *xCoM* crossing the border of support. One way to do that is to adjust the ipsilateral *CoP* position (the ipsilateral border of support) by placing the ipsilateral forepaw more laterally or medially depending on a *xCoM* deviation. Note that the ipsilateral hindpaw is already on the ground during the diagonal double support phase and cannot be used to adjust the *CoP* position (**Figure 2B**, mouse).

We quantified animal responses to *xCoM* deviations (Δ*xCoM*) by determining the deviation of *CoP* position (black continuous circles in **Figure 2B**) from the mean ipsilateral *CoP* position computed across all strides within a given trial (black dashed circles). Another potential mechanism for adjusting lateral balance is quickening or delaying the initiation of the ipsilateral double support phase by lifting the contralateral forelimb. This creates the moment of gravitational force in the frontal plane applied to the animal body that can be considered an inverted pendulum in this phase (Latash et al., 2020). This gravitational moment decelerates *CoM* motion towards the ipsilateral border of support and then reverses it. We considered the elapsed time between the middle of the diagonal (or quadrupedal) support phase and onset of the ipsilateral double support phase (*t_r_*, **Figure 2B**) as the second characteristic by which the animal could regulate lateral balance.

### Potential effects of instrumentation on locomotion

Implantation of EMG electrodes in both cats and mice, as well as the placement of reflective markers on cats, could potentially alter locomotor behavior. For example, during early stages of training while wearing reflective markers, some cats adopted a more crouched walking posture. To assess whether instrumentation affected locomotor stability measures, we compared step width and margin of stability (*MoS*) before and after EMG electrode implantation in additional animals.

Specifically, step width and *MoS* were measured in two additional mice walking on a treadmill at 0.1 m/s and in two cats from the present study walking overground at self-selected speeds. For the cats, walking speeds before and after implantation were 0.82 ± 0.07 m/s and 0.70 ± 0.07 m/s for cat 1, and 0.82 ± 0.04 m/s and 0.63 ± 0.05 m/s for cat 2, respectively.

In cats, hindlimb step width decreased following implantation in both tested animals. In cat 1, hindlimb step width decreased from 5.1 ± 0.9 cm to 4.0 ± 0.9 cm (22%; number of steps 𝑛_1_ = 36, 𝑛_2_ = 20; *p* < 0.001, independent-samples *t*-test), and in cat 2 from 3.7 ± 0.9 cm to 3.3 ± 1.1 cm (11%; 𝑛_1_ = 47, 𝑛_2_ = 65; *p* = 0.018). The right-side *MoS* decreased in cat 1 following implantation (3.4 ± 0.5 cm vs. 2.3 ± 0.6 cm; 32%; number of cycles 𝑛_1_ = 17, 𝑛_2_ = 12; *p* < 0.001) but did not change significantly in cat 2 (1.8 ± 0.5 cm vs. 1.9 ± 0.7 cm; 𝑛_1_ = 26, 𝑛_2_ = 32; *p* = 0.561).

In mice, hindlimb step width increased following implantation in one animal, rising from 2.1 ± 0.2 cm to 2.4 ± 0.2 cm in mouse 2 (13%; 𝑛_1_ = 40, 𝑛_2_ = 42; *p* < 0.05), whereas no significant change was observed in mouse 1 (2.2 ± 0.3 cm vs. 2.3 ± 0.3 cm; 𝑛_1_ = 36, 𝑛_2_ = 46; *p* = 0.190). In contrast to step width, both mice exhibited significant increases in right-side *MoS* following implantation: from 0.9 ± 0.2 cm to 1.1 ± 0.8 cm in mouse 1 (20%; 𝑛_1_ = 27, 𝑛_2_ = 32; *p* < 0.05) and from 0.8 ± 0.2 cm to 1.1 ± 0.9 cm in mouse 2 (25%; 𝑛_1_ = 18, 𝑛_2_ = 40; *p* < 0.05).

As discussed later, these differences were relatively small and did not affect the main conclusions of the study.

### Statistics

To compare major kinematic and lateral stability variables between cats and mice, we used linear mixed-effects models (MIXED, IBM SPSS Statistics v29, USA). Models included either a single fixed factor, *Species* (mouse, cat), or two fixed factors, *Species* and *Joint* (ankle, knee, hip), as appropriate. Post hoc comparisons were conducted using Bonferroni-corrected multiple comparisons.

To determine whether or how the animals regulate their lateral balance in each step, we used a linear regression equation in the form 𝑌 = 𝑏_0_ + 𝑏_1_𝑋 to predict animal response, a dependent variable 𝑌 (*t_r_* or Δ𝐶𝑜𝑃) as a function of the *CoM* motion state *X* (Δ𝑥𝐶𝑜𝑀); coefficients 𝑏_0_and 𝑏_1_ are regression coefficients. High and significant correlation coefficient between the dependent and independent variables would support the idea of within step regulation of lateral balance.

For all statistical tests, the significance level was set to 𝑝 = 0.05.

## RESULTS

### General kinematic characteristics of locomotion in cats and mice

At a treadmill speed of 0.4 m/s, cats locomoted exclusively using a walking gait, as indicated by the limb support diagram (thick black horizontal bars in **Figure 2B** showing stance phases of each limb) and by the absence of an aerial phase. The mean stride cycle duration was 0.878 ± 0.138 s (mean ± SD), and the normalized, dimensionless cycle duration (𝑡^_𝑐_) was 5.91 ± 0.91 (**Figure 3A, B**). Mean stance duration was 0.654 ± 0.121 s (**Figure 3A**), and the duty factor of the right forelimb exceeded 0.5 (0.742 ± 0.065; **Figure 3C**), consistent with a walking gait (Hildebrand, 1989).

**Figure 3.**
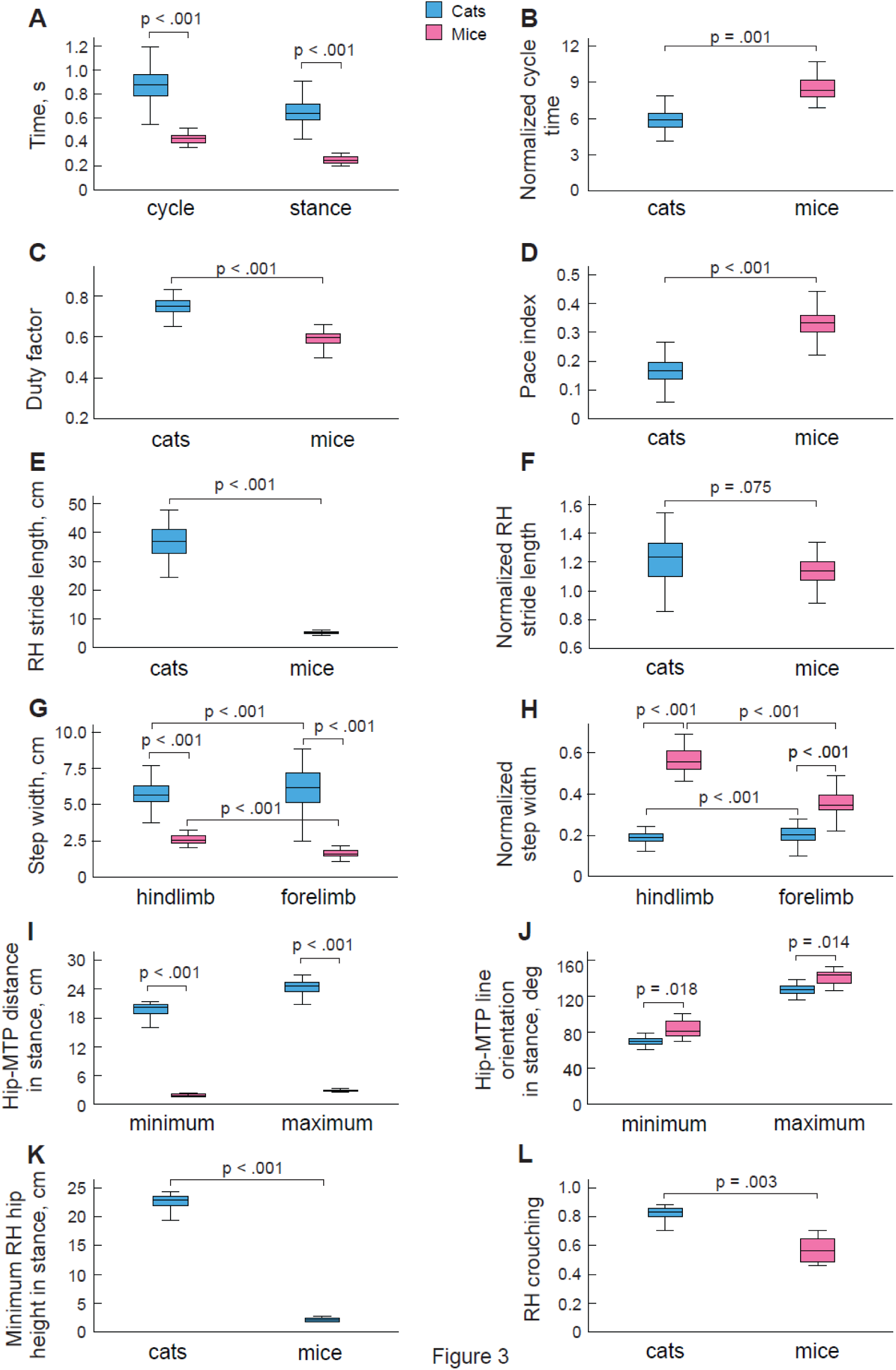
Boxplots of general and right hindlimb kinematic characteristics of locomotion of cats and mice. P-values indicate statistical significance (linear mixed effect model analysis, total number of cycles across 4 cats is n=128 and across 5 mice, n=58). **A**: Cycle time and stance time. Both variables have significantly greater values for cats. **B**: Normalized cycle time, 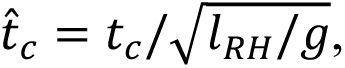 where 𝑡_𝑐_ is the cycle time in seconds, 𝑙_𝑅𝐻_ is the total hindlimb length (the sum of lengths of the thigh, shank, and tarsals; Table 1), and 𝑔 is gravitational acceleration. The normalized cycle time is significantly greater in mice. **C**: Duty cycle. It is significantly greater in cats. **D**: Pace index. It is significantly lower in cats indicating that movement phases of their ipsilateral limbs are closer to each other. **E**, **F**: Right hindlimb (RH) absolute and normalized stride length, respectively. RH stride length was normalized by 𝑙_𝑅𝐻_. The absolute RH stride length is significantly greater in cats; there is no difference in normalized RH stride length between cats and mice. **G**, **H**: Hindlimb step width, absolute and normalized by 𝑙_𝑅𝐻_, respectively, for hindlimbs and forelimbs. The absolute values of step width for hindlimbs and forelimbs are significantly greater in cats. In cats, absolute and normalized forelimb step widths are significantly greater than those of hindlimb step width (related-samples Wilcoxon signed rank test); in mice, the forelimb step width is significantly smaller than the hindlimb step width. **I, J**: The minimum and maximum RH length (the hip-MTP distance, 𝑙_ℎ𝑖𝑝−𝑀𝑇𝑃_) in cm and normalized by 𝑙_𝑅𝐻_, respectively, in stance. The absolute minimum and maximum RH lengths are significantly greater in cats; the normalized RH lengths are significantly greater in mice. **K**: Minimum RH hip height in stance. It is significantly greater in cats. **L**: RH crouching index in stance. The maximum value of crouching index of 1 indicates full limb extension. Mice have significantly smaller crouching index, i.e. greater crouching.

At a treadmill speed of 0.1 m/s, mice frequently trotted but occasionally walked; to enable direct comparison between species, only walking trials were analyzed. The mean duty factor for mice was 0.592 ± 0.034 (**Figure 3C**), confirming a walking gait. Mean cycle and stance durations were 0.427 ± 0.041 s and 0.253 ± 0.030 s, respectively, and the normalized cycle duration (𝑡^_𝑐_) was 8.57 ± 0.95 (**Figure 3A–C**). Absolute values of cycle duration, stance duration, and duty factor were significantly greater in cats than in mice (F₁,₇ = 55.1–133.6, all *p* < 0.001). In contrast, the normalized cycle duration (𝑡^_𝑐_) was significantly greater in mice than in cats (F₁,₇ = 34.3, *p* < 0.001), reflecting animal size-dependent differences in gait timing.

The pattern of limb support during walking differed between species, resembling pacing in cats and trotting in mice. The pace index—defined as the phase difference between ipsilateral hindlimb and forelimb footfalls—ranges from ∼0 to 0.05 for pacing and from ∼0.45 to 0.55 for trotting (Hildebrand, 1989). Consistent with these definitions, the pace index was significantly lower in cats than in mice (0.173 ± 0.091 vs. 0.333 ± 0.048; F₁,₇ = 132.8, *p* < 0.001; **Figure 3D**).

Right hindlimb stride length in cats was approximately sevenfold greater than in mice (36.8 ± 5.15 cm vs. 5.13 ± 0.62 cm; F₁,₇ = 359.8, *p* < 0.001; **Figure 3E**). Normalization of stride length by each animal’s right hindlimb anatomical length (𝑙_RH_) eliminated this difference (1.217 ± 0.154 in cats vs. 1.125 ± 0.142 in mice; F₁,₇ = 4.67, *p* = 0.075; **Figure 3F**). In contrast, normalization by the mean hip–metatarsophalangeal distance during stance (𝑙_hip-MTP_) revealed significantly shorter normalized strides in cats than in mice (1.702 ± 0.236 vs. 2.110 ± 0.316; F₁,₇ = 7.75, *p* = 0.033), reflecting limb posture-dependent difference in stride length.

During the swing phase, both forepaws and hindpaws of cats moved laterally along a bell-shaped trajectory with symmetric rising and falling slopes and a peak near mid-swing (**Figure 2B**). During stance, both forepaws and hindpaws were placed closer to the body midline relative to their swing-phase positions. On average, the lateral distance between the left and right forepaws at stance onset (forelimb stance width) in cats was greater than the corresponding hindlimb stance width (6.15 ± 1.37 cm vs. 5.71 ± 0.82 cm; *p* < 0.001, Wilcoxon signed-rank test; **Figure 3G**).

In contrast, mice exhibited much smaller mediolateral paw excursions during swing, with forepaws and hindpaws following relatively straight trajectories (**Figure 2B**). In mice, forelimb stance width was significantly smaller than hindlimb stance width (1.65 ± 0.33 cm vs. 2.59 ± 0.29 cm; *p* < 0.001, Wilcoxon signed-rank test; **Figure 3G**). After normalization of both forelimb and hindlimb step widths by right anatomical hindlimb length (𝑙_RH_), normalized step widths were significantly greater in mice than in cats (F₁,₇ = 55.4–300.1, *p* < 0.001; **Figure 3H**).

Figure 2A illustrates representative examples of hindlimb posture during the stance phase in a cat and a mouse, highlighting the orientation of the thigh, shank, and tarsal segments. Qualitatively, the most prominent postural difference between species was shank orientation: in mice, the shank was oriented closer to horizontal. Together with greater joint flexion, this posture resulted in a lower hip height and a shorter hip–metatarsophalangeal (MTP) distance 𝑙_hip-MTP_ during stance in mice.

Mean hip height during the stance phase was 22.2 ± 1.9 cm in cats and 2.2 ± 0.3 cm in mice (F₁,₇ = 382.4, *p* < 0.001; Figure 3K). The minimum 𝑙_hip-MTP_ distance during stance was 19.3 ± 2.3 cm in cats and 1.9 ± 0.3 cm in mice (F₁,₇ = 160.0, *p* < 0.001; Figure 3I), while the corresponding maximum values during stance were 23.9 ± 2.1 cm and 2.9 ± 0.2 cm, respectively (F₁,₇ = 296.8, *p* < 0.001; Figure 3I). During stance, right-hindlimb orientation varied between minimum angles of 68.8° ± 5.9° in cats and 83.4° ± 8.7° in mice, and maximum angles of 126.7° ± 5.5° and 141.3° ± 7.6°, respectively (Figure 3J). Both minimum and maximum orientation angles were significantly greater in mice than in cats (F₁,₇ = 10.4–11.8, *p* = 0.014–0.018).

To quantify limb crouching, we normalized the minimum hip height during stance by the anatomical hindlimb length (𝑙_RH_; **Table 1**). This ratio was significantly greater in cats than in mice (0.812 ± 0.068 vs. 0.572 ± 0.080; F₁,₇ = 19.3, *p* = 0.003; Figure 3L), indicating a more crouched hindlimb posture in mice (smaller ratio corresponds to greater crouching) (Biewener, 1983; Day and Jayne, 2007). The absolute and normalized hip height values observed here are consistent with previously reported measurements in cats and mice during locomotion (Trank et al., 1996; Day and Jayne, 2007; Vemula et al., 2019).

### Joint angles

Consistent with the more crouched posture of mice, hindlimb joint angles were substantially more flexed throughout both stance and swing in mice compared to cats, although the overall temporal patterns of joint motion were similar across species (Figure 4). The large difference in the absolute magnitude of hip joint angles between cats and mice was partly attributable to differences in hip angle definition (Figure 2A). The time course of right-hindlimb length (hip–metatarsophalangeal distance, 𝑙_hip-MTP_) closely resembled the knee angle pattern, particularly in cats, whereas the time course of hindlimb orientation more closely matched the hip angle pattern (Figure 4).

**Figure 4.**
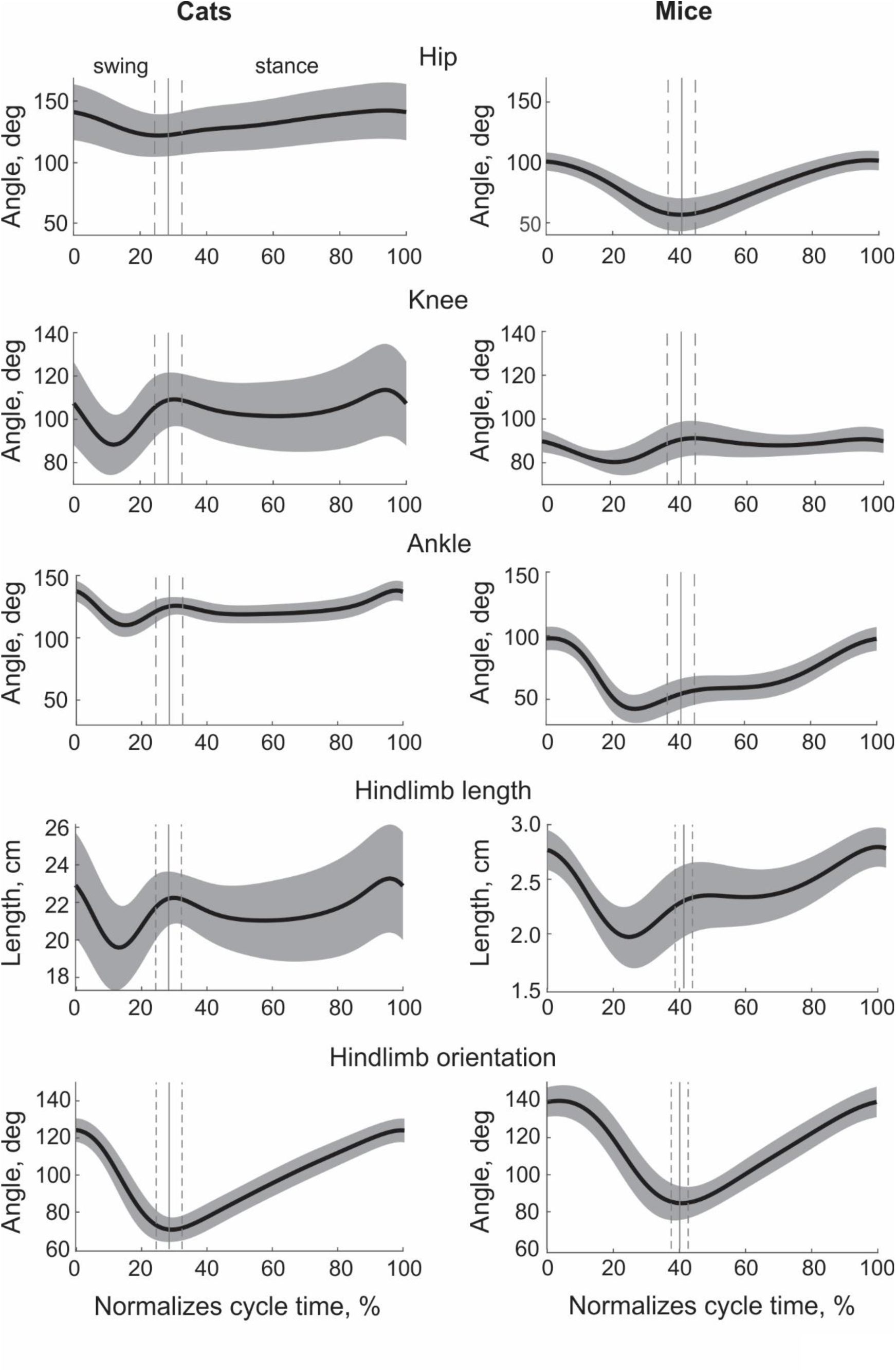
Right hindlimb kinematic variables as a function of the normalized cycle time during walking of cats and mice. The solid thick lines are the means averaged across all subjects and walking cycles within cats and mice; gray areas are standard deviations (SD). The total number of cycles across 4 cats is n=128 and across 5 mice, n=58. Vertical continuous vertical line and thin dashed lines in each panel indicate the mean normalized time and SD of stance onset, respectively. The panels from top to bottom are hip joint angles, knee joint angles, ankle joint angles, hindlimb length (𝑙_ℎ𝑖𝑝−𝑀𝑇𝑃_), and hindlimb orientation (see Figure 2 for definitions of the latter two variables).

Consistent with a more extended hindlimb posture in cats, minimum and maximum joint angles during stance and swing were generally greater in cats than in mice, except for knee joint angles. Minimum knee angles during stance were comparable between cats and mice (98.9 ± 16.8° vs. 85.2 ± 4.6°; F₁,₇ = 2.8, *p* = 0.133), as were minimum knee angles during swing (86.9 ± 14.2° vs. 79.3 ± 5.2°; F₁,₇ = 0.215, *p* = 0.656; Figure 5A**, C**). During stance, minimum ankle angles were 116.7 ± 7.8° in cats and 52.9 ± 9.7° in mice (F₁,₇ = 97.1, *p* < 0.001), and minimum hip angles were 122.0 ± 17.0° and 56.0 ± 13.4°, respectively (F₁,₇ = 106.5, *p* < 0.001; Figure 5A).

**Figure 5.**
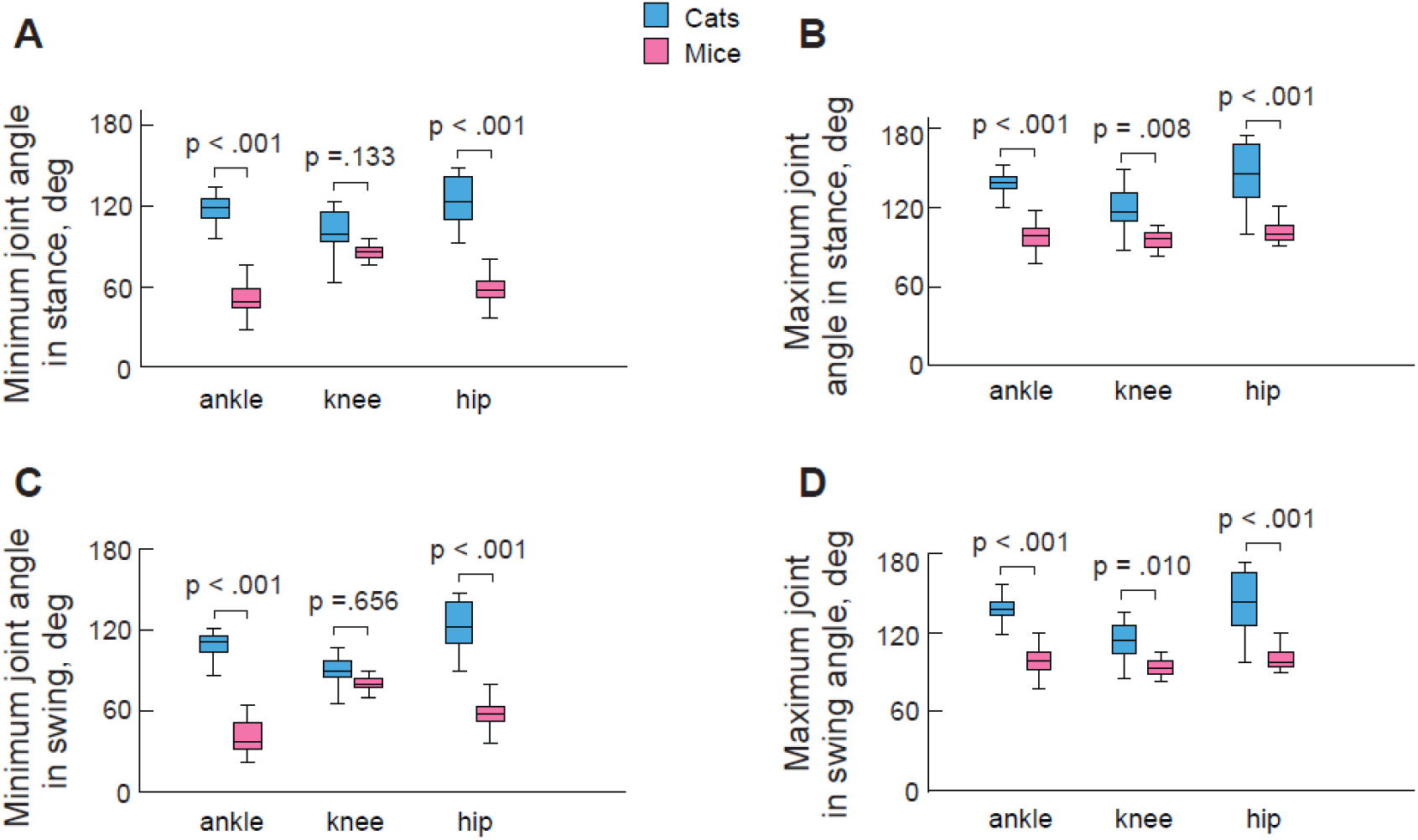
Boxplots of minimum and maximum joint angles of right hindlimb joints during the stance and swing phases of walking cats and mice. P-values indicate statistical significance (linear mixed effect model analysis, total number of cycles across 4 cats is n=128 and across 5 mice, n=58). **A**: Minimum ankle, knee, and hip joint angles in stance. **B**: Maximum ankle, knee, and hip joint angles in stance. **C**: Minimum ankle, knee, and hip joint angles in swing. **D**: Maximum ankle, knee, and hip joint angles in swing.

Maximum joint angles during stance were also greater in cats than in mice. Maximum ankle angles were 138.8 ± 7.5° in cats and 97.5 ± 9.3° in mice (F₁,₇ = 43.1, *p* < 0.001), maximum knee angles were 118.5 ± 15.1° and 95.7 ± 6.1° (F₁,₇ = 11.9, *p* = 0.008), and maximum hip angles were 143.4 ± 23.4° and 101.9 ± 8.0°, respectively (F₁,₇ = 43.9, *p* < 0.001; Figure 5B).

During swing, minimum ankle angles were 116.7 ± 7.8° in cats and 52.9 ± 9.7° in mice (F₁,₇ = 86.2, *p* < 0.001), and minimum hip angles were 121.5 ± 17.1° and 56.0 ± 13.3°, respectively (F₁,₇ = 82.3, *p* < 0.001; Figure 5C). Maximum swing-phase joint angles were greater in cats than in mice for the ankle (137.9 ± 7.7° vs. 98.3 ± 9.1°; F₁,₇ = 48.3, *p* < 0.001), knee (114.2 ± 12.5° vs. 94.2 ± 6.3°; F₁,₇ = 11.2, *p* = 0.010), and hip (141.9 ± 23.2° vs. 101.9 ± 8.0°; F₁,₇ = 52.2, *p* < 0.001; Figure 5D).

Overall, the joint angle patterns and the minimum and maximum joint angle values observed in cats and mice are consistent with previously reported measurements during locomotion (Lavoie et al., 1995; Carlson-Kuhta et al., 1998; Leblond et al., 2003; Akay et al., 2014; Charles et al., 2018; Gregor et al., 2018).

### Kinematics of lateral *CoM* motion

During walking in both cats and mice, lateral positions of the *CoM* and *xCoM* oscillated periodically within the extreme left and right positions of the center of pressure (*CoP*), which correspond to ipsilateral double-support phases (Figure 2B). A walking cycle was defined as the interval between right forelimb (RF) swing onset and the subsequent RF swing onset (shaded rectangles in Figure 2B). Within each cycle, two ipsilateral double-support periods occurred, during which the body was supported by either the two left or the two right limbs. The extreme left and right *CoP* positions therefore corresponded to the lateral boundaries of the support area, and their separation approximated the average of the hindlimb and forelimb step widths (Figures 2B**, 3G, H**). Body stability is compromised when the *CoM* moves beyond the boundary of support (Hof et al., 2005; Hof et al., 2007).

The temporal patterns of *CoM* and *xCoM* lateral displacements were similar to those reported previously for cats (Park et al., 2019; Latash et al., 2020) and humans (Hof et al., 2007; Buurke et al., 2019) walking on split-belt or tied-belt treadmills. Specifically, *xCoM* exhibited larger peak excursions than *CoM* and reached these peaks slightly earlier, near the onset of ipsilateral double support (Figure 2B). The lateral *CoM* velocity waveform was phase-shifted by approximately one-quarter of the cycle relative to the *CoM* displacement. *CoM* velocity peaks occurred between successive *CoM* peaks around the middle of the diagonal double support periods (or quadrupedal support periods) (Figure 2B, cat; dashed vertical lines).

The mean lateral displacement of the *CoM* between leftward and rightward peaks was significantly greater in cats than in mice (2.4 ± 0.5 cm vs. 0.45 ± 0.17 cm; F₁,₇ = 169.5, *p* < 0.001; Figure 6A). When normalized by anatomical hindlimb length (𝑙_RH_), this difference was no longer significant (0.078 ± 0.019 vs. 0.098 ± 0.036; F₁,₇ = 3.77, *p* = 0.092; Figure 6B). Similarly, the absolute range of lateral *CoM* velocity was significantly larger in cats than in mice (14.4 ± 2.7 cm/s vs. 5.4 ± 1.7 cm/s; F₁,₇ = 86.8, *p* < 0.001; Figure 6C), whereas no significant interspecies difference was observed when velocity range was normalized using the Froude number (0.099 ± 0.020 vs. 0.110 ± 0.032; F₁,₇ = 0.405, *p* = 0.545; Figure 6D).

**Figure 6.**
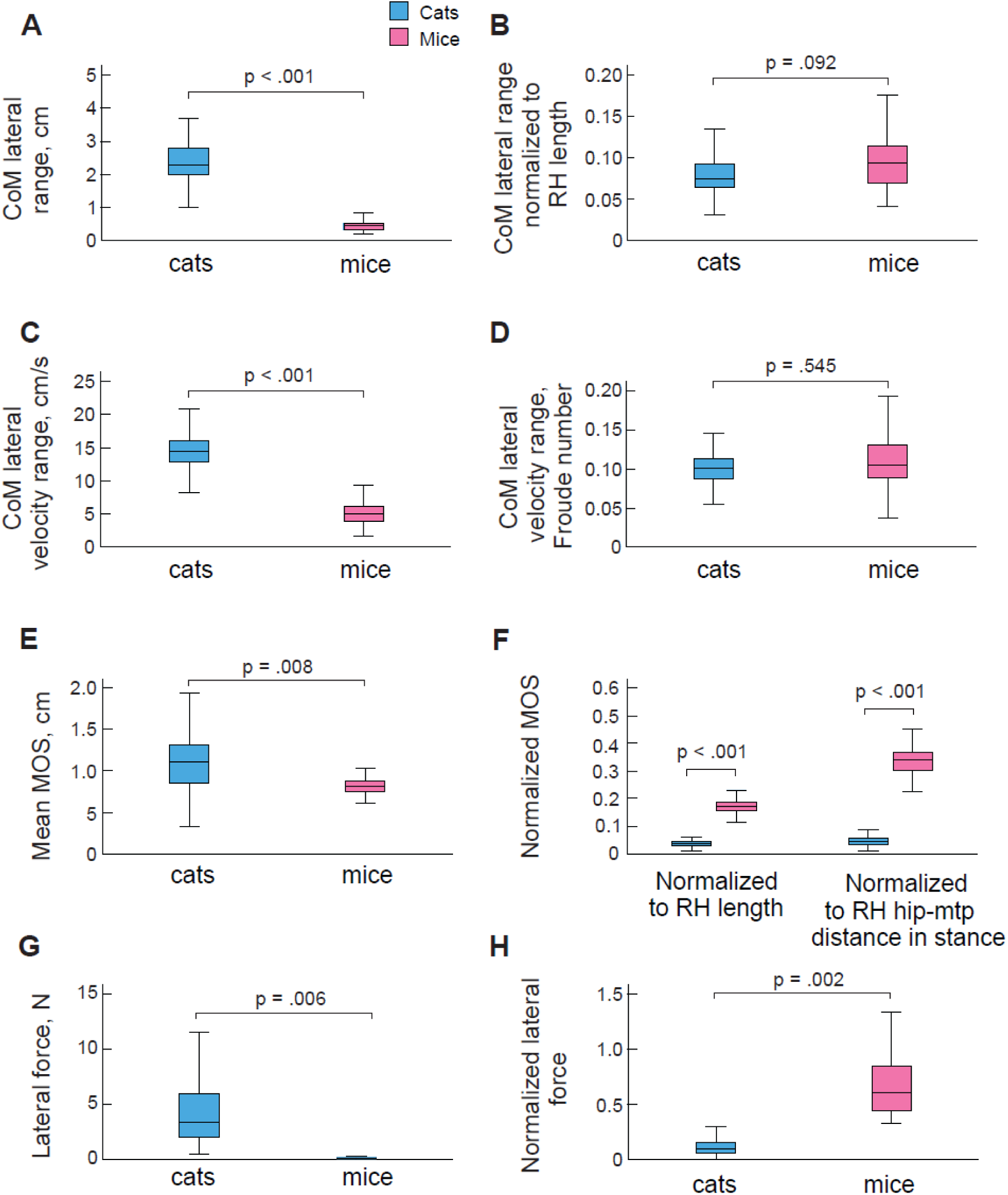
Boxplots of characteristics of lateral *CoM* motion and body dynamic stability during walking of cats and mice. P-values indicate statistical significance (linear mixed effect model analysis, total number of cycles across 4 cats is n=128 and across 5 mice, n=58). **A, B**: Absolute and normalized by hindlimb length (𝑙_𝑅𝐻_) range of lateral *CoM* motion, respectively. The absolute *CoM* range is significantly greater in cats, but there is no significant difference in the normalized *CoM* range between cats and mice. **C, D**: Absolute and normalized (Froude number, see Methods) range of lateral *CoM* velocity, respectively. The absolute *CoM* velocity range is significantly greater in cats, but there is no significant difference in the normalized *CoM* velocity range between cats and mice. **E, F**: Absolute and normalized by the hindlimb length (𝑙_𝑅𝐻_) and hip-MTP distance (𝑙_ℎ𝑖𝑝−𝑀𝑇𝑃_) margins of dynamic stability (*MoS*), respectively. Note that *MoS*/𝑙_𝑜_ (where 𝑙_𝑜_ is a scaling factor proportional to subject’s linear size) corresponds to the normalized (dimensionless) impulse of force 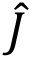 that causes *xCoM* to cross the boundary of support when applied to *CoM* at the instant of *xCoM* peak: 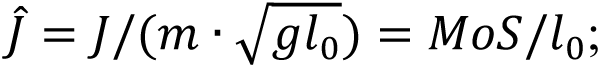 see text and (Nguyen et al., 2023). The absolute MoS is significantly greater in cats than in mice. Normalization by 𝑙_𝑅𝐻_ and 𝑙_ℎ𝑖𝑝−𝑀𝑇𝑃_make the normalized *MoS* significantly greater in mice than in cats. **G**, **H**: The absolute and normalized by body weight, respectively, lateral force applied to CoM during the ipsilateral double-support phase that causes the body to lose balance (see text for more details). The absolute destabilizing force is statistically greater in cats; the normalized force is statistically greater in mice.

### Measures of lateral stability

We compared lateral dynamic stability between cats and mice using the margin of stability (*MoS*). In absolute terms, *MoS* was significantly greater in cats than in mice (1.10 ± 0.33 cm vs. 0.82 ± 0.13 cm; F₁,₇ = 12.5, *p* = 0.008; Figure 6E). However, when *MoS* was normalized by anatomical hindlimb length (𝑙_RH_) or by the mean hip–metatarsophalangeal distance during stance (𝑙_hip-MTP_), cats exhibited markedly smaller values than mice (0.036 ± 0.011 vs. 0.180 ± 0.027; F₁,₄ = 403.2, *p* < 0.001, and 0.051 ± 0.017 vs. 0.338 ± 0.066; F₁,₄ = 279.2, *p* < 0.001, respectively; Figure 6F). Thus, when accounting for body size and limb posture, relative lateral dynamic stability was substantially lower in cats than in mice.

A consistent pattern emerged when stability was quantified using an alternative measure: the lateral force required to destabilize the body during the ipsilateral double-support phase (𝐹_lat_). The absolute magnitude of this force was significantly greater in cats than in mice (4.6 ± 3.7 N vs. 0.151 ± 0.070 N; F₁,₈ = 13.6, *p* = 0.006; Figure 6G). However, when normalized by body weight, the destabilizing force required for cats was much smaller than for mice: approximately 13% of body weight in cats versus 68% in mice (0.134 ± 0.097 vs. 0.678 ± 0.285; F₁,₆ = 31.7, *p* = 0.002; Figure 6H).

### Initiation of ipsilateral double-support period and adjustment of *CoP* in response to *xCoM* deviation from the midline

In search for potential mechanisms regulating lateral dynamic stability, we computed regression equations and correlations between the independent variable characterizing the state of *CoM* lateral motion, Δx𝐶𝑜𝑀, and the dependent variables which represent a potential response of the control system to potential destabilization of lateral balance, *t_r_* and Δ𝐶𝑜𝑃 (see Methods and Figure 2B).

There was no significant relationship between the elapsed time *t_r_* and Δx𝐶𝑜𝑀 in cats (r = 0.093, F_1,264_ = 2.307, p = 0. 130) and mice (r = 0.153, F_1,106_ = 2.543, p = 0. 114); Figure 7A**, B**. On the other hand, the Δ𝐶𝑜𝑃 was moderately correlated with the Δx𝐶𝑜𝑀 in both cats (r = 0.639; F_1,276_ = 190.0, p < 0. 001) and mice (r = 0.616, F_1,106_ = 64.9, p < 0. 001); Figure 7 **C, D**. Thus, the Δx𝐶𝑜𝑀 explain ∼40% of Δ𝐶𝑜𝑃 variance in cats (r^2^ = 0.408) and mice (r^2^ = 0.380) supporting the notion of step-by-step control of lateral stability in both species.

**Figure 7.**
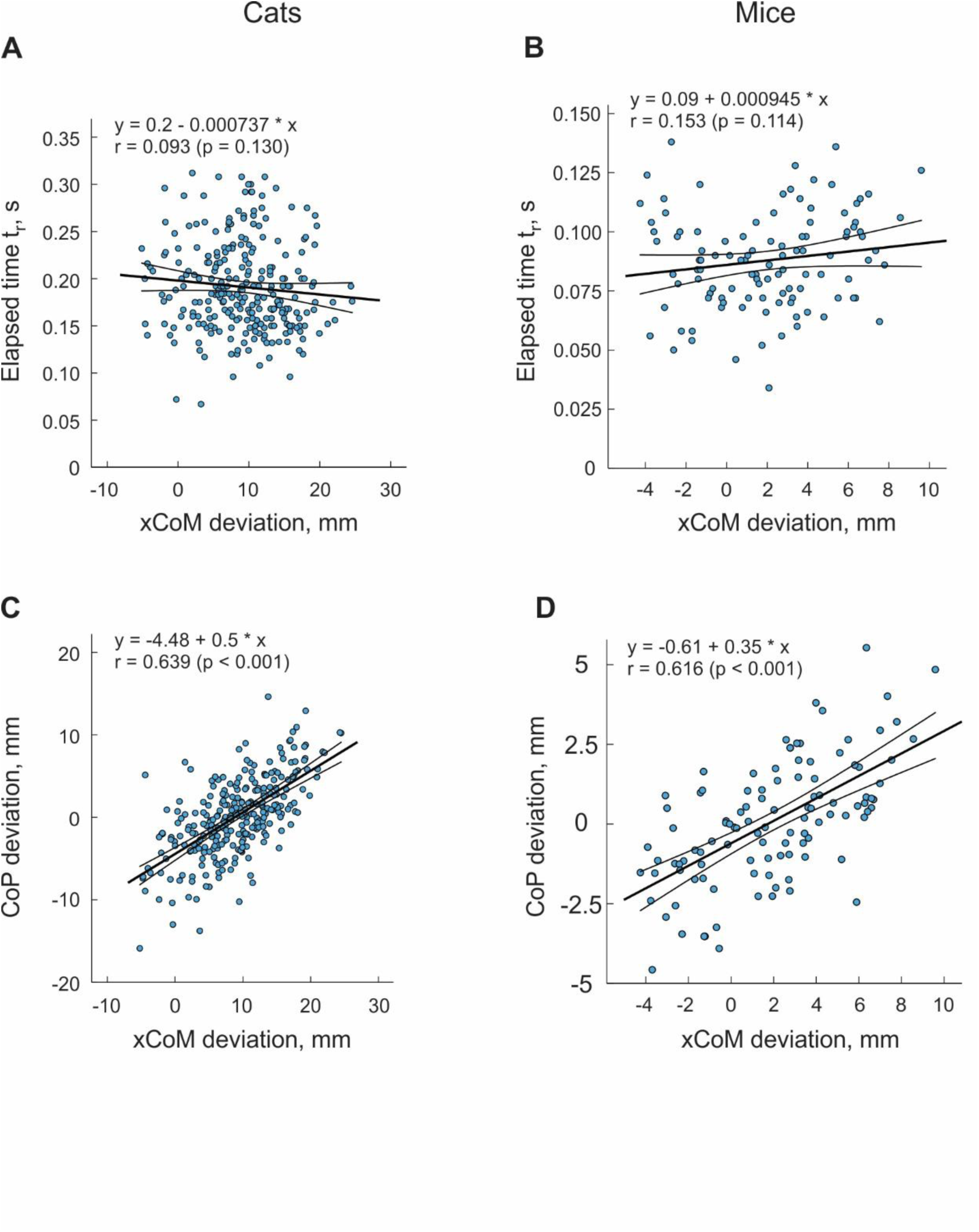
Results of regression and correlation analyses between lateral deviations of the extrapolated center of mass (*xCoM*), characterizing the state of lateral *CoM* motion, and two kinematic variables considered as potential corrective responses of the animal (see text for more details). The blue dots represent experimental points for all walking cycles of cat (n = 266) and mice (n =108). The regression line (± mean 95% confidence interval) describes the linear fit in the form 𝑦 = 𝑏_0_ + 𝑏_1_𝑥, where *y* and *x* are the variables plotted along the vertical and horizontal axes, and *b_0_* and *b_1_* are regression coefficients; *r* is the Pearson correlation coefficient. **A, B**: Scatter plots of the relationship between the elapsed time *t_r_* and the *xCoM* deviation from the mean for cats and mice. The relationship is not significant for cats (F_1,264_ = 2.307, p = 0. 130) and mice (F_1,106_ = 2.543, p = 0. 114). **C, D**: Scatter plots of the relationship between the *CoP* deviation from the mean and the *xCoM* deviation from the mean for cats and mice. The relationship is significant for cats (F_1,276_ = 190.0, p < 0. 001) and mice (F_1,106_ = 64.9, p < 0. 001).

## DISCUSSION

### Posture and support geometry shape lateral stability

We compared lateral dynamic stability between two quadrupedal mammals that differ markedly in body size and limb posture and found that, when expressed in size-normalized terms, stability is substantially greater in mice than in cats. This difference reflects the combined effects of relatively wider step width and more crouched limb posture in mice, rather than effects of body size.

Lateral stability during locomotion depends on both support geometry and *CoM* dynamics. Although the absolute margin of dynamic stability (*MoS*) was larger in cats, normalization by anatomical hindlimb length or posture-related distance 𝑙_ℎ𝑖𝑝−𝑀𝑇𝑃_revealed markedly greater stability in mice. These findings highlight a key limitation of displacement-based metrics of postural stability when comparing across species of different sizes and confirm the utility of dimensionless measures that incorporate both body dimensions and dynamics (Hof et al., 2005; Nguyen et al., 2023).

The enhanced stability of mice is driven primarily by differences in support geometry and limb posture. Mice exhibit significantly wider relative step widths, which expand the mediolateral base of support. In contrast, normalized excursions of the *xCoM* were similar between species, indicating that differences in stability arise largely from how the support base is configured rather than from differences in *CoM* motion itself. Normalization of *MoS* by posture-dependent distance 𝑙_ℎ𝑖𝑝−𝑀𝑇𝑃_further amplified interspecies differences, underscoring the contribution of crouching. A lower hip height reduces the effective inverted-pendulum length and increases margin of stability for a given *CoM* displacement (Alexander, 2002). Together, these findings support the long-standing hypothesis that crouched postures enhance locomotor stability (Walter, 2003; Blum et al., 2011).

### Mechanical versus neural contributions to stability

The relationship between body size and stability reflects competing mechanical and neural factors. Larger animals benefit from greater inertia, which resists external perturbations, but they also face longer sensorimotor delays relative to stride duration. Conversely, smaller animals experience reduced inertial stability but compensate through posture and support geometry. Consistent with this trade-off, absolute lateral forces required to destabilize the body were greater in cats, yet when normalized to body weight, mice required substantially larger perturbations. These results indicate that relative stability is greater in mice even though larger animals are mechanically more resistant to perturbations in absolute terms (Mohamed Thangal and Donelan, 2020).

These findings suggest a shift in the balance between mechanical and neural stabilization strategies across body sizes. Mice, with their wider stance and crouched posture, may rely more heavily on passive mechanical stability somewhat similar to that in cockroaches (Full et al., 2002), whereas cats may depend more on active neural control to maintain balance. This interpretation is consistent with prior work showing that rodents retain locomotor stability despite substantial sensory or spinal lesions, whereas cats exhibit impaired balance under comparable conditions (Stapley et al., 2002; Fouad et al., 2010; Akay et al., 2014; Audet et al., 2023).

### Step-by-step control of lateral balance

Despite differences in overall stability, both species exhibited evidence of step-by-step control of lateral balance. Specifically, deviations of the extrapolated *CoM* (*xCoM*) were correlated with subsequent adjustments of the center of pressure (*CoP*) at the ipsilateral double support phase, indicating that animals modulate the support base to counteract destabilizing motion. Unlike bipedal locomotion, in which lateral stability is primarily regulated through step placement, quadrupedal walking limits the effectiveness of forelimb lateral placement because the ipsilateral hindlimb is already in stance. This constraint is particularly pronounced in mice, where forelimb step width is smaller than hindlimb step width, reducing the capacity of the forelimbs to modulate the mediolateral base of support. In contrast, cats exhibit greater forelimb than hindlimb step width, a difference that may reflect feline functional and morphological specialization of the forelimbs for balance control (Hudson et al., 2011; Rahmati et al., 2025).

### Implications for evolution of locomotor function

Our findings identify limb posture and support geometry as primary determinants of lateral stability across mammals. Smaller animals adopt crouched postures and relatively wider step widths, which enhance stability but increase energetic costs and reduce locomotor speed. In contrast, larger animals achieve greater locomotor efficiency and speed and reduced musculoskeletal stress by adopting more extended limb postures, often at the expense of stability (Taylor et al., 1982; Biewener, 1989). These trade-offs have important evolutionary implications. In small mammals, enhanced stability may arise as a byproduct of evolutionary pressures favoring a low body profile and increased maneuverability, particularly because stability and falls are less consequential than in larger species (Hooper, 2012). A lower body height, combined with greater maneuverability, facilitates movement through cluttered environments, enabling rapid changes in direction and improving the likelihood of escaping predators by accessing narrow refuges. In contrast, larger mammals may rely more heavily on high-speed or long-distance locomotion to find food and escape predators at the expense of active sensorimotor control to maintain balance and prevent falls.

### Relevance for neural control and translational research

Differences in posture-dependent stability have important implications for interpreting animal models of locomotion. Rodents are widely used in neuroscience because of genetic accessibility, yet their crouched posture and high intrinsic stability may reduce reliance on neural control mechanisms that are critical in larger animals including humans (Duysens et al., 2000; Frigon et al., 2021).

Accordingly, caution is warranted when extrapolating findings from rodent models to human balance control. Cats, which exhibit lower relative stability and greater dependence on sensory feedback, may provide a closer analogue for studying neural mechanisms of locomotor balance and their disruption in disease. Notably, the dimensionless measures of lateral locomotor stability observed in cats, e.g., *MoS*/𝑙_𝑅𝐻_= 0.04, are of the same order of magnitude as those reported in humans, 0.02-0.03 (Nguyen et al., 2023), suggesting that larger mammals, including bipeds, may share common constraints and control strategies.

This similarity supports the idea that findings in cats may generalize more readily to human balance control than those in small rodents.

### Study limitations

Several potential limitations of the present study should be noted. First, the method used to estimate *CoM* displacements in mice based on a ventral marker may be less accurate than volume-based body reconstructions. However, direct validation against a body-volume–based approach showed that mean differences in estimated *CoM* displacements were relatively small, remaining below 10%. Second, implantation of EMG electrodes caused measurable changes in step width and margin of stability (*MoS*) in individual animals, including a 20–25% increase in *MoS* in mice and a 32% decrease in one cat. Importantly, these effects were severalfold smaller than the interspecies differences in normalized *MoS* observed between mice and cats. Therefore, neither potential inaccuracies in *CoM* estimation nor implantation-related changes in locomotor kinematics are likely to have influenced the main conclusions of this study.

Another potential limitation is that the walking speeds of mice (0.1 m/s) and cats (0.4 m/s) did not correspond to identical Froude numbers (see Methods) and therefore may not reflect perfectly dynamically similar locomotion (Alexander and Jayes, 1983). Notably, 0.1 m/s is close to the upper end of the self-selected walking speed range in mice (Lemieux et al., 2016; Machado et al., 2020), whereas 0.4 m/s is near the lower end of the self-selected walking speed range in cats (Bishop et al., 2008; Beloozerova et al., 2010). Although it would have been possible to analyze faster mouse locomotion to better match the Froude number used for cats, this would likely have required comparison of cat walking with mouse trotting. Such a comparison would introduce a qualitatively different gait pattern and, consequently, greater dynamic dissimilarity between species. Therefore, we prioritized gait-matched comparisons over exact Froude-number matching to ensure meaningful assessment of lateral stability mechanisms during walking in both species.

## Conclusion

Lateral dynamic stability during quadrupedal locomotion depends strongly on limb posture and support geometry rather than body size alone. Smaller mammals achieve greater relative stability through crouched postures and wider steps, whereas larger mammals operate closer to stability limits and rely more on active control. These findings identify limb posture and support geometry as key factors shaping locomotor evolution and provide a framework for linking biomechanics with neural control across species of different sizes.

## ACKNOWLEDGMENTS

We thank Brenda Ross for technical assistance. We also thank Rachel Watermeier and Lamisa Rahman for her help with data analysis.

## REFERENCES

Akay T, Acharya HJ, Fouad K, Pearson KG (2006) Behavioral and electromyographic characterization of mice lacking EphA4 receptors. J Neurophysiol 96:642–651.

Akay T, Tourtellotte WG, Arber S, Jessell TM (2014) Degradation of mouse locomotor pattern in the absence of proprioceptive sensory feedback. Proceedings of the National Academy of Sciences of the United States of America 111:16877–16882.

Alexander RM (2002) Stability and manoeuvrability of terrestrial vertebrates. Integr Comp Biol 42:158–164.

Alexander RMN, Jayes AS (1983) A dynamic similarity hypothesis for the gaits of quadrupedal mammals. J Zool 201:135–152.

Audet J, Yassine S, Lecomte CG, Mari S, Soucy F, Morency C, Merlet AN, Harnie J, Beaulieu C, Gendron L, Rybak IA, Prilutsky BI, Frigon A (2023) Spinal Sensorimotor Circuits Play a Prominent Role in Hindlimb Locomotor Recovery after Staggered Thoracic Lateral Hemisections but Cannot Restore Posture and Interlimb Coordination during Quadrupedal Locomotion in Adult Cats. eNeuro 10.

Bauby CE, Kuo AD (2000) Active control of lateral balance in human walking. J Biomech 33:1433–1440.

Beloozerova IN, Farrell BJ, Sirota MG, Prilutsky BI (2010) Differences in movement mechanics, electromyographic, and motor cortex activity between accurate and nonaccurate stepping. J Neurophysiol 103:2285–2300.

Biewener AA (1983) Allometry of quadrupedal locomotion: the scaling of duty factor, bone curvature and limb orientation to body size. J Exp Biol 105:147–171.

Biewener AA (1989) Scaling body support in mammals: limb posture and muscle mechanics. Science 245:45–48.

Bishop KL, Pai AK, Schmitt D (2008) Whole body mechanics of stealthy walking in cats. PloS one 3:e3808.

Blum Y, Birn-Jeffery A, Daley MA, Seyfarth A (2011) Does a crouched leg posture enhance running stability and robustness? J Theor Biol 281:97–106.

Buurke TJW, Lamoth CJC, van der Woude LHV, Hof AL, den Otter R (2019) Bilateral temporal control determines mediolateral margins of stability in symmetric and asymmetric human walking. Sci Rep 9:12494.

Carlson-Kuhta P, Trank TV, Smith JL (1998) Forms of forward quadrupedal locomotion. II. A comparison of posture, hindlimb kinematics, and motor patterns for upslope and level walking. J Neurophysiol 79:1687–1701.

Charles JP, Cappellari O, Hutchinson JR (2018) A Dynamic Simulation of Musculoskeletal Function in the Mouse Hindlimb During Trotting Locomotion. Front Bioeng Biotechnol 6:61.

Day LM, Jayne BC (2007) Interspecific scaling of the morphology and posture of the limbs during the locomotion of cats (Felidae). J Exp Biol 210:642–654.

De Comite A, Seethapathi N (2025) Foot placement control underlies stable locomotion across species. Proceedings of the National Academy of Sciences of the United States of America 122:e2413958122.

Dick TJ, Clemente CJ (2017) Where Have All the Giants Gone? How Animals Deal with the Problem of Size. PLoS Biol 15:e2000473.

Duysens J, Clarac F, Cruse H (2000) Load-regulating mechanisms in gait and posture: comparative aspects. Physiol Rev 80:83–133.

Farrell BJ, Bulgakova MA, Beloozerova IN, Sirota MG, Prilutsky BI (2014) Body stability and muscle and motor cortex activity during walking with wide stance. Journal of neurophysiology 112:504–524.

Fischer MS, Blickhan R (2006) The tri-segmented limbs of therian mammals: kinematics, dynamics, and self-stabilization--a review. Journal of experimental zoology Part A, Comparative experimental biology 305:935–952.

Fouad K, Rank MM, Vavrek R, Murray KC, Sanelli L, Bennett DJ (2010) Locomotion after spinal cord injury depends on constitutive activity in serotonin receptors. J Neurophysiol 104:2975–2984.

Frigon A (2020) Fundamental contributions of the cat model to the neural control of locomotion. In: The Neural Control of Movement (Whelan PJ, Sharples SA, eds), pp 315–348. London, UK: Academic Press.

Frigon A, Akay T, Prilutsky BI (2021) Control of mammalian locomotion by somatosensory feedback. Compr Physiol 12:2877–2947.

Fuentes MA (2016) Theoretical considerations on maximum running speeds for large and small animals. J Theor Biol 390:127–135.

Full RJ, Kubow T, Schmitt J, Holmes P, Koditschek D (2002) Quantifying dynamic stability and maneuverability in legged locomotion. Integr Comp Biol 42:149–157.

Gambaryan PP (1974) How Animals Run: Anatomical Adaptations. New York: John Wiley and Sons.

Gates DH, Scott SJ, Wilken JM, Dingwell JB (2013) Frontal plane dynamic margins of stability in individuals with and without transtibial amputation walking on a loose rock surface. Gait Posture 38:570–575.

Gatesy SM, Biewener AA (1991) Bipedal locomotion: effects of speed, size and limb posture in birds and humans. J Zool 224:127–147.

Goslow GE, Jr., Reinking RM, Stuart DG (1973) The cat step cycle: hind limb joint angles and muscle lengths during unrestrained locomotion. J Morphol 141:1–41.

Gray J (1968) Animal locomotion. London: William Clowes & Sons.

Gregor RJ, Maas H, Bulgakova MA, Oliver A, English AW, Prilutsky BI (2018) Time course of functional recovery during the first 3 mo after surgical transection and repair of nerves to the feline soleus and lateral gastrocnemius muscles. J Neurophysiol 119:1166–1185.

Hildebrand M (1989) The quadupedal gaits in vertebrates. BioScience 39:766–775.

Hof AL (2018) Scaling and Normalization. In: Handbook of Human Motion (Müller B, Wolf SI, eds), pp 295–305: Springer Nature.

Hof AL, Gazendam MG, Sinke WE (2005) The condition for dynamic stability. J Biomech 38:1–8.

Hof AL, van Bockel RM, Schoppen T, Postema K (2007) Control of lateral balance in walking. Experimental findings in normal subjects and above-knee amputees. Gait Posture 25:250–258.

Hooper SL (2012) Body size and the neural control of movement. Current biology: CB 22:R318–322.

Horner AM, Hanna JB, Biknevicius AR (2016) Crouching to fit in: the energetic cost of locomotion in tunnels. J Exp Biol 219:3420–3427.

Hoy MG, Zernicke RF (1985) Modulation of limb dynamics in the swing phase of locomotion. J Biomech 18:49–60.

Hudson PE, Corr SA, Payne-Davis RC, Clancy SN, Lane E, Wilson AM (2011) Functional anatomy of the cheetah (Acinonyx jubatus) forelimb. J Anat 218:375–385.

Klishko AN, Akyildiz A, Mehta-Desai R, Prilutsky BI (2021) Common and distinct muscle synergies during level and slope locomotion in the cat. J Neurophysiol 126:493–515.

Klishko AN, Farrell BJ, Beloozerova IN, Latash ML, Prilutsky BI (2014) Stabilization of cat paw trajectory during locomotion. J Neurophysiol 112:1376–1391.

Latash EM, Barnett WH, Park H, Rider JM, Klishko AN, Prilutsky BI, Molkov YI (2020) Frontal plane dynamics of the centre of mass during quadrupedal locomotion on a split-belt treadmill. J R Soc Interface 17:20200547.

Lavoie S, McFadyen B, Drew T (1995) A kinematic and kinetic analysis of locomotion during voluntary gait modification in the cat. Exp Brain Res 106:39–56.

Leblond H, L’Esperance M, Orsal D, Rossignol S (2003) Treadmill locomotion in the intact and spinal mouse. The Journal of neuroscience: the official journal of the Society for Neuroscience 23:11411–11419.

Lemieux M, Josset N, Roussel M, Couraud S, Bretzner F (2016) Speed-Dependent Modulation of the Locomotor Behavior in Adult Mice Reveals Attractor and Transitional Gaits. Frontiers in neuroscience 10:42.

Machado AS, Marques HG, Duarte DF, Darmohray DM, Carey MR (2020) Shared and specific signatures of locomotor ataxia in mutant mice. Elife 9.

Mathis A, Mamidanna P, Cury KM, Abe T, Murthy VN, Mathis MW, Bethge M (2018) DeepLabCut: markerless pose estimation of user-defined body parts with deep learning. Nat Neurosci 21:1281–1289.

Modi AD, Parekh A, Patel ZH (2024) Methods for evaluating gait associated dynamic balance and coordination in rodents. Behavioural brain research 456:114695.

Mohamed Thangal SN, Donelan JM (2020) Scaling of inertial delays in terrestrial mammals. PloS one 15:e0217188.

Molina LK, Small GH, Neptune RR (2023) The influence of step width on balance control and response strategies during perturbed walking in healthy young adults. J Biomech 157:111731.

More HL, Donelan JM (2018) Scaling of sensorimotor delays in terrestrial mammals. Proceedings Biological sciences / The Royal Society 285.

Nevo E (1979) Adaptive Convergence and Divergence of Subterranean Mammals. Annual Review of Ecology and Systematics 10:269–308.

Nguyen NT, Christensen MS, Tracy JB, Kellaher GK, Pohlig RT, Crenshaw JR (2023) How should the margin of stability during walking be expressed to account for body size? J Biomech 161:111835.

Pantall A, Gregor RJ, Prilutsky BI (2012) Stance and swing phase detection during level and slope walking in the cat: Effects of slope, injury, subject and kinematic detection method. J Biomech 45:1529–1533.

Park H, Latash EM, Molkov YI, Klishko AN, Frigon A, DeWeerth SP, Prilutsky BI (2019) Cutaneous sensory feedback from paw pads affects lateral balance control during split-belt locomotion in the cat. J Exp Biol 222.

Pearson KG, Acharya H, Fouad K (2005) A new electrode configuration for recording electromyographic activity in behaving mice. J Neurosci Methods 148:36–42.

Percie du Sert N et al. (2020) The ARRIVE guidelines 2.0: Updated guidelines for reporting animal research. PLoS Biol 18:e3000410.

Prilutsky BI, Parker J, Cymbalyuk GS, Klishko AN (2022) Emergence of Extreme Paw Accelerations During Cat Paw Shaking: Interactions of Spinal Central Pattern Generator, Hindlimb Mechanics and Muscle Length-Depended Feedback. Frontiers in integrative neuroscience 16:810139.

Prilutsky BI, Maas H, Bulgakova M, Hodson-Tole EF, Gregor RJ (2011) Short-term motor compensations to denervation of feline soleus and lateral gastrocnemius result in preservation of ankle mechanical output during locomotion. Cells Tissues Organs 193:310–324.

Rahmati SM, Klishko AN, Martin RS, Bunderson NE, Meslie JA, Nichols TR, Rybak IA, Frigon A, Burkholder TJ, Prilutsky BI (2025) Role of forelimb morphology in muscle sensorimotor functions during locomotion in the cat. J Physiol 603:447–487.

Rankin BL, Buffo SK, Dean JC (2014) A neuromechanical strategy for mediolateral foot placement in walking humans. J Neurophysiol 112:374–383.

Riskin DK, Kendall CJ, Hermanson JW (2016) The crouching of the shrew: Mechanical consequences of limb posture in small mammals. PeerJ 4:e2131.

Santuz A, Laflamme OD, Akay T (2022) The brain integrates proprioceptive information to ensure robust locomotion. J Physiol 600:5267–5294.

Stapley PJ, Ting LH, Hulliger M, Macpherson JM (2002) Automatic postural responses are delayed by pyridoxine-induced somatosensory loss. The Journal of neuroscience: the official journal of the Society for Neuroscience 22:5803–5807.

Taylor CR, Heglund NC, Maloiy GM (1982) Energetics and mechanics of terrestrial locomotion. I. Metabolic energy consumption as a function of speed and body size in birds and mammals. J Exp Biol 97:1–21.

Thirkell JE, Bennett NC, Hart DW, Faulkes CG, Daley MA, Portugal SJ (2025) Metabolic expenditure of submaximal locomotion in naked mole-rats (Heterocephalus glaber) and Damaraland mole-rats (Fukomys damarensis). J Exp Biol 228.

Townsend MA (1985) Biped gait stabilization via foot placement. J Biomech 18:21–38.

Trank TV, Chen C, Smith JL (1996) Forms of forward quadrupedal locomotion. I. A comparison of posture, hindlimb kinematics, and motor patterns for normal and crouched walking. J Neurophysiol 76:2316–2326.

Vemula MG, Deliagina TG, Zelenin PV (2019) Kinematics of forward and backward locomotion performed in different environmental conditions. J Neurophysiol 122:2142–2155.

Vidal PP, Degallaix L, Josset P, Gasc JP, Cullen KE (2004) Postural and locomotor control in normal and vestibularly deficient mice. J Physiol 559:625–638.

Walter RM (2003) Kinematics of 90 degrees running turns in wild mice. J Exp Biol 206:1739–1749.

Wang Y, Srinivasan M (2014) Stepping in the direction of the fall: the next foot placement can be predicted from current upper body state in steady-state walking. Biology letters 10.

